# Oncofetal protein CRIPTO regulates wound healing and fibrogenesis in regenerating liver and is associated with the initial stages of cardiac fibrosis

**DOI:** 10.1101/2021.06.29.450195

**Authors:** S. Karkampouna, D. van der Helm, B. van Hoek, H.W. Verspaget, M.J. Goumans, M.J. Coenraad, B.P.T. Kruithof, M. Kruithof-de Julio

**Affiliations:** Department for Biomedical Research, Urology Research, Bern University, Bern, Switzerland; Department of Gastroenterology and Hepatology, Leiden University Medical Center, Leiden, the Netherlands; Department of Cell and Chemical Biology, Leiden University Medical Center, Leiden, the Netherlands; Department of Cardiology, Leiden University Medical Center, Leiden, the Netherlands; Department of Urology, Inselspital, Bern University Hospital, Bern, Switzerland

**Author notes:** **Corresponding Author:** Prof. Dr. M. Kruithof-de Julio, Department for Biomedical Research, Urology Research, University of Bern, Bern, Switzerland, Murtenstrasse 35, 3008. equal contribution (S. Karkampouna, D. van der Helm). Joint senior authorship (M.J. Coenraad, B.P.T. Kruithof, M. Kruithof-de Julio).

## Abstract

**Background:** Oncofetal protein, Cripto, is silenced during postnatal life and often re-expressed in different neoplastic processes. In the present study we investigated the potential role of Cripto in hepatic and cardiac fibrosis. In this study, the aim was to explore whether Cripto is expressed during liver fibrogenesis and whether this is related to the disease severity and pathogenesis of fibrogenesis. Furthermore, we aimed to identify the impact of Cripto expression on fibrogenesis in organs with high versus low regenerative capacity, represented by murine liver fibrogenesis and adult murine heart fibrogenesis

**Methods:** Circulating CRIPTO levels were measured in plasma samples of patients with cirrhosis registered at the waitlist for liver transplantation (LT) and one year after LT. The expression of Cripto and fibrotic markers (aSMA, collagen I) were determined in human liver tissues of patients with cirrhosis (on a basis of viral hepatitis or alcoholic disease), in cardiac tissue samples of patients with end-stage heart failure and of mice with experimental liver and heart fibrosis using immuno-histochemical stainings and qPCR. Mouse models with experimental chronic liver fibrosis, induced with multiple shots of carbon tetrachloride (CCl_4_) and acute liver fibrosis (one shot of CCl_4_) were evaluated for Cripto expression and fibrotic markers. Cripto was overexpressed *in vivo* (Adenoviral delivery) or functionally sequestered by ALK4Fc ligand trap in the acute liver fibrosis mouse model. Murine heart tissues were evaluated for Cripto and fibrotic markers, in three models of heart injury; following myocardial infarction, pressure overload and *ex vivo* induced fibrosis.

**Results:** Patients with end-stage liver cirrhosis showed elevated Cripto levels in plasma, which had decreased one year after LT. Cripto expression was observed in fibrotic tissues of patients with end-stage liver cirrhosis and in patients with heart failure. The expression of Cripto in the liver was found specifically in the hepatocytes and was positively correlated with the Model for End-stage Liver Disease (MELD) score for end-stage liver disease. Cripto expression in the samples of cardiac fibrosis was limited and mostly observed in the interstitial cells. In the chronic and acute mouse models of liver fibrosis, Cripto-positve cells were observed in damaged liver areas around the central vein, which preceded the expression of aSMA-positive stellate cells, i.e. mediators of fibrosis. Whereas in the chronic mouse models the fibrosis and Cripto expression was still present after 11 weeks, in the acute model the liver regenerated and the fibrosis and Cripto expression resolved. *In vivo* overexpression of Cripto in this model, led to an increase in fibrotic markers while blockage of Cripto secreted function inhibited the extend of fibrotic areas and marker expression (αSMA, Collagen type I and III) and induced higher proliferation of residual healthy hepatocytes. Cripto expression was also upregulated in several mouse models of cardiac fibrosis. During myocardial infarction Cripto is upregulated initially in cardiac interstitial cells, followed by expression in αSMA-positive myofibroblasts throughout the infarct area. After the scar formation, Cripto expression decreased concomitantly with the aSMA expression. Temporal expression of Cripto in αSMA-positive myofibroblasts was also observed surrounding the coronary arteries in the pressure overload model of cardiac fibrosis. Furthermore, Cripto expression was upregulated in interstitial myofibroblasts in hearts cultured in an ex vivo model for cardiac fibrosis.

**Conclusion:** Our results are indicative for a functional role of Cripto in induction of fibrogenesis and potential applications in antifibrotic treatments and stimulation of tissue regeneration.

## Introduction

Fibrotic diseases are responsible for 45% of death in the developed world [1], therefore understanding the regulatory pathways involved in organ fibrosis is a necessary step. Fibrosis is the increased secretion and deposition of extracellular matrix (ECM) that will lead to perturbation of the normal tissue architecture and eventually results in organ dysfunction [2, 3]. In some organs/tissues the fibrotic tissue can be (partly) resolved by regenerative processes as seen in liver fibrosis. In other organs, the fibrosis is permanent due to the inabilitly of the organ to regenerate, as seen in the heart. Whereas fibrosis can occur in many different organs/tissues and involves the interplay of many different cell types and factors, the fibrotic process in the different organs/tissus share many similarities including their triggers, the activation of myofibroblasts (MFBs), involvement of inflammation and the activation of the Transforming Growth Factor-beta (TGF-β) pathway [1].

ECM-producing MFBs are the key mediators of tissue remodelling and characterized by aSMA expression [4–6]. The source of the MFBs is organ dependent [7]. For example, in the liver they derive from the hepatic stellate cells, i.e. pericytes found in the space of Disse, whereas in the heart they derive from the existing interstitial fibroblasts [8, 9]. The activation of the MFBs is often the result of chronic exposure to damaging factors.

In the liver, the stellate cells are thought to be activated by damaged and apoptotic hepatocytes upon chronic exposure to damaging factors such as alcohol and viral hepatitis B or C (HBV, HCV) [10]. Approximately 10-30% of chronic liver diseases progresses to cirrhosis, which is associated with high mortality and health care burden [11]. The excessive collagen deposition disrupts liver architecture, leading to hepatocellular dysfunction [2] [12–14] ultimately cirrhosis and eventually increases the risk of hepatocellular carcinoma (HCC) development.

During myocardial infarction (MI), cardiomyocyte death leads to the generation of many inflammatory and profibrotic factors that can activate the interstitial fibroblasts. The MFBs subsequently form a fibrous scar, which is critical to protect the heart from rupture. Hypertension or aortic stenosis, on the other hand, causes increased ventricular pressure, leading to the activation of MFBs and subsequent formation of progressive interstitial and perivascular fibrosis [15]. The fibrosis in the heart triggers myocardial stiffness, resulting in ventricular dysfunction, which could ultimately result in heart failure [3].

Hepatic fibrosis, advanced cirrhosis, and cardiac fibrosis are major health problems, with a lack of effective antifibrotic treatment options [16–19]. For liver fibrosis, withdrawal of the injuring stimulus is the only current treatment for liver fibrosis, which in some cases leads to the resolution of fibrogenesis[20]. For end-stage liver cirrhosis, liver transplantation (LT) is still the only curative treatment option of which feasibility depends on patient condition and donor availability, but LT is still a major surgical intervention with substantial risk of complications and risk of disease recurrence [21, 22]. For cardiac fibrosis current therapies aim to promote more functional scar formation and to improve heart function [23], however, effective therapies directly targeting the process of fibrosis do not exist. Therefore, therapies directly targeting fibrosis are needed. Better understanding of the pathological mechanisms underlying fibrosis could lead to identification of new biomarkers to monitor the disease and may also lead to new targets for the development of alternative treatment strategies.

Cripto is a GPI-anchored signalling protein, member of the epidermal growth factor-Cripto/frl/cryptic (EGF-CFC) family, with diverse functions in embryogenesis and as regulator of stemness [24, 25]. Cripto is silenced postnatally and often re-expressed in neoplasms of breast, lung, prostate, ovarian, bladder, colon, skin, and brain, were it is thought to be involved in cancer progression and metastasis [26–35]. Recently, we observed Cripto expression in the majority of cirrhotic liver tissue [36], which might indicate that reactivation of Cripto in adult tissues is associated with pathological conditions such as inflammation, fibrosis and (pre)malignant state. Therefore, we assessed in the present study whether Cripto is expressed during fibrosis and whether this was related to the disease severity. To determine whether Cripto expression in fibrosis is liver-specific or rather a general feature of fibrosis, we expanded our study to heart fibrosis.

Cripto expression was evaluated in liver cirrhosis specimens and in circulating blood levels of patients with cirrhosis prior to and after removal of the fibrogenic liver by LT. A correlation of Cripto expression with disease stage could imply a functional role for Cripto in the fibrosis–cirrhosis-HCC cascade rendering it a potential interesting marker for disease monitoring or even as a treatment target. Cripto expression was further evaluated in validated mouse models for acute and chronic liver fibrosis; chronic or single administration of hepatotoxin carbon tetrachloride (CCl_4_), respectively. In three models of heart injury; following myocardial infarction, pressure overload and *ex vivo* induced fibrosis, Cripto was found to be temporally upregulated. Our findings indicate that Cripto is an immediate wound healing response gene, following tissue injury of different aetiologies and is a master orchestrator of fibrogenesis in multiple tissues as determined in human cirrhosis, murine hepatic models, as well as *in vivo* and *ex vivo* cardiac fibrosis. Our data imply Cripto as a potential marker for disease monitoring and a treatment target of organ fibrosis.

## Materials and Methods

### Patients and controls

Plasma Cripto levels were measured in paired pre- and 1 year post-LT plasma samples from consecutive patients with end-stage liver disease due to alcoholic cirrhosis (ALD, N=25) or viral hepatitis-induced cirrhosis (N=20) who had a plasma sample available available from just before transplantation. Cripto levels measured in plasma from healthy volunteers (N=16) served as control. Exclusion criteria for this study were the presence of HCC, a combined etiology of cirrhosis, death or re-LT within one year after LT and the development of serious adverse events after LT, such as Tacrolimus induced renal insufficiency (Supplemental table 1: patient characteristics). For qPCR and (immuno)-histochemical analysis, control tissue (N=5) and alcohol- or viral hepatitis-induced fibrotic/cirrhotic liver tissue (N=19) were obtained from the tissue collections of the LUMC Liver diseases Biobank and Pathology department. These tissues were derived from different patients than the patients included in the circulating plasma Cripto study. All liver tissues were obtained during LT, or resection of HCC or colorectal cancer-derived liver metastasis.

Clinical data were extracted from electronic patient files, including laboratory assessments and clinical MELD (Model for End-Stage Liver Disease) scores, a scoring system for assessing liver function impairment in cirrhosis and risk of short-term mortality. Cardiac samples were resected from the left ventricle of patients with end-stage heart failure (Kindly provided by Dr. Bax). All experiments with human specimens were approved by the ethical research committee of the Leiden University Medical Center (LUMC, protocol number: B15.006). Materials were used in compliance with the rules prescribed by the regulations of the LUMC Liver diseases Biobank and with a signed informed consent of the donors.

### Mouse models of fibrosis

All animal experiments were performed in C57BL6 mice in compliance with the guidelines for animal care and approved by the LUMC Animal Care Committee. Mice received food and water *ad libitum* and were housed under 12h day/night cycle. Liver fibrosis was induced in 6 week old male mice as described previously [37]. For a period of 11 weeks, mice received 2 intraperitoneal (i.p) injections per week with carbon tetrachloride (CCl_4_) in mineral oil (Sigma-Aldrich Chemie BV, Zwijndrecht, The Netherlands). The first week, mice received 2 initiating higher dosages of CCl_4_ of 1 ml/kg. The following 10 weeks a maintenance dose of 0.75 ml/kg was given twice weekly. At the end of 11^th^ week, mice were sacrificed and livers collected and subsequently fixated with 4% paraformaldehyde (PFA) for paraffin embedding or stored in isobutyl for RNA isolation.

Acute liver injury was induced in 5-6 weeks old male C57Bl6 mice weighing 20-25 g by i.p. injection of a single dose of 1 ml/kg body weight carbon tetrachloride (CCL_4_) (Sigma), in mineral oil or mineral oil alone as control. Mice were sacrificed after 3, 6, 24, 48, 72 hours and day 6 (n=2-3 per time point).

Sections of mouse hearts with induced MI or TAC were kindly provided by Dr. Smits. MI was induced in 10–12 weeks old mice by ligation of the left anterior descending artery (LAD), as described previously [38]. Hearts were collected at 1, 3, 7, 14 and 28 days post-MI. Pressure overload of the left ventricle was induced by transverse aortic constriction, (TAC), which was performed as described previously [39]. Hearts were isolated 2, 6, and 8 weeks post-TAC. *Ex vivo* culturing of adult mouse hearts was performed as described [40, 41] using the ligation of the aorta and induction of retrograde flow of 1000 μl/min for 7 days. Hearts were fixed overnight in 4% PFA/PBS at 4 °C.

### *In vivo* administration of adenovirus

Adenoviral constructs expressing β-galactosidase (lacZ) or Cripto were prepared using the Gateway adenoviral expression vectors pAd/CMV/V5-lacZ or pAd/CMV/V5-DEST, mouse Cripto expression plasmid was a gift from from Dr. Peter Gray [42]. AdlacZ (AdCon) or AdCripto (1×10E+9 viral particles/ mouse) was injected intravenously via the tail vein to enhance delivery to the liver [43]. After 24 hours, CCl_4_ was injected i.p. (day 0). Treatment groups were as follows: AdlacZ, CCl_4_+AdlacZ, AdCripto, CCl_4_+AdCripto. At day 1, day 2 and day 3 after CCl_4_ administration mice were sacrificed and liver tissues were collected for histology preparation, RNA/ protein isolation and analysis.

### Histological examination of fibrosis

Tissues fixed in PFA were dehydrated through a graded series of ethanol, cleared in xylene, embedded in paraffin, and sectioned at 4-6 μm. To evaluate the severity of fibrosis in human and mouse liver tissue, a Sirius-red staining was performed to visualize the amount of collagen deposition. Paraffin sections (4 μm) were hydrated and subsequently stained for 90 min with 1 g/L Sirius-red F3B in saturated picric acid (both Klinipath). Next, the sections were incubated for 10 min with 0.01 M HCL, dehydrated and mounted with Entellan (Merck KGaA, Darmstadt, Germany). Fixed microscope settings were used to capture 5-8 representative images (10x magnifications) which were subsequently used to quantify the amount of staining with ImageJ software (ImageJ 1.47v, National Institutes of Health, USA). With fixed threshold settings, based on control tissues, positive pixels were measured and the respective percentage to the total image calculated and defined as the positive area.

### Immunohistochemistry, immunofluorescence imaging and quantification

Immunohistochemical stainings were performed in fibrotic and control human and mouse liver tissue. Briefly, paraffin tissue sections (4 μm) were hydrated and endogenous peroxidases blocked with 0.3% H_2_O_2_/methanol (20 min). Antigen retrieval was performed by 10 min boiling in citrate buffer (0.1 M, pH 6.0). After cooling down, primary antibodies detecting mouse- and human-Cripto (both kindly provided by Dr Gray Clayton Foundation Laboratories for Peptide Biology, The Salk Institute for Biological Studies, La Jolla, California, USA) and anti-αSMA antibodies (A2547, clone 1A_4_; Sigma, Buchs, Switzerland) were added and incubated overnight. Human-αSMA staining was visualised by 1h incubation with a secondary goat anti-rabbit-HRP conjugated antibody followed by a 10 min incubation with 3,3’-diaminobenzidine (DAB Fast Tablet, Sigma-Aldrich, St. Louis, MO). Nuclear counterstaining was performed with hematoxylin after which the sections were dehydrated and mounted with Entellan.

For immunofluorescence for the acute liver and heart models of fibrosis, tissues were fixed overnight in 4% PFA/PBS and paraffin sections were sectioned at 4 μm (liver tissues) or 6 μm (heart tissues). Antigen retrieval was performed in pressure cooker (30min liver, 40min heart tissues) in Antigen Unmasking citrate-based solution (H-3300) Vector laboratories, USA) and sections were blocked in 1% BSA solution/0.01%Tween20 in PBS. The following primary antibodies were added: Cripto and αSMA, anti-Collagen type I (1:500, 1310-01, Southern Biotech, USA) and anti-Collagen type III (1:500, 1330-01, Southern Biotech, USA), anti-Ki67 (1:200, clone SP6, GTX16667, GeneTex, LucernaChem, Switzerland). The following day, sections were incubated with secondary antibodies labelled with Alexa Fluor 488, 555, or 647 (Invitrogen/Molecular Probes, Zurich, Switzerland; 1:250 in PBS/0.1% Tween 20). Tyramide Signal Amplification (PerkinElmer) was used to amplify the Cripto signal followed by Alexa Fluor® 488 streptavidin (Invitrogen) for visualization as previously described[36]. Sections were counterstained with TO-PRO-3 (Invitrogen/Molecular Probes) or DAPI solution (Sigma) for visualization of nuclei, and mounted using Prolong G anti-fade mounting medium (Invitrogen/Molecular Probes). Representative pictures of the liver samples were captured using a confocal microscope (Leica Biosystems BV, Amsterdam, The Netherlands) and 40x 1.4NA oil-immersion objective with fixed microscope and software settings. Subsequently, 5-10 representative pictures were captured and used for quantification. The amount of DAB or fluorescent staining in the representative pictures was quantified with ImageJ software. For the Cripto stainings in Fig. 1 and Fig. 2 the threshold was set, based on control tissues, and defined as a percentage of positive pixels compared to the total pixels within the hepatocyte regions, whereas regions such as the vessels, bile ducts and the septa with ECM were excluded from the analysis. For the αSMA stainings, whole images were used to quantify the positive area. For the immunofluorescence quantification of Cripto, Collagen type I, III and αSMA staining, the positive area was measured with fixed threshold settings and represent as percentage. For the Ki67 quantification, the number of positive cells was normalized to total number of nuclei (DAPI) on each section and an average of three different fields of view areas was used for each liver tissue. All slides with cardiac samples were scanned with the Pannoramic 250 slide scanner (version1.23, 3DHISTECH Ltd.) and analyzed using Caseviewer (version2.3, 3DHISTECH Ltd.).

**Figure 1.**
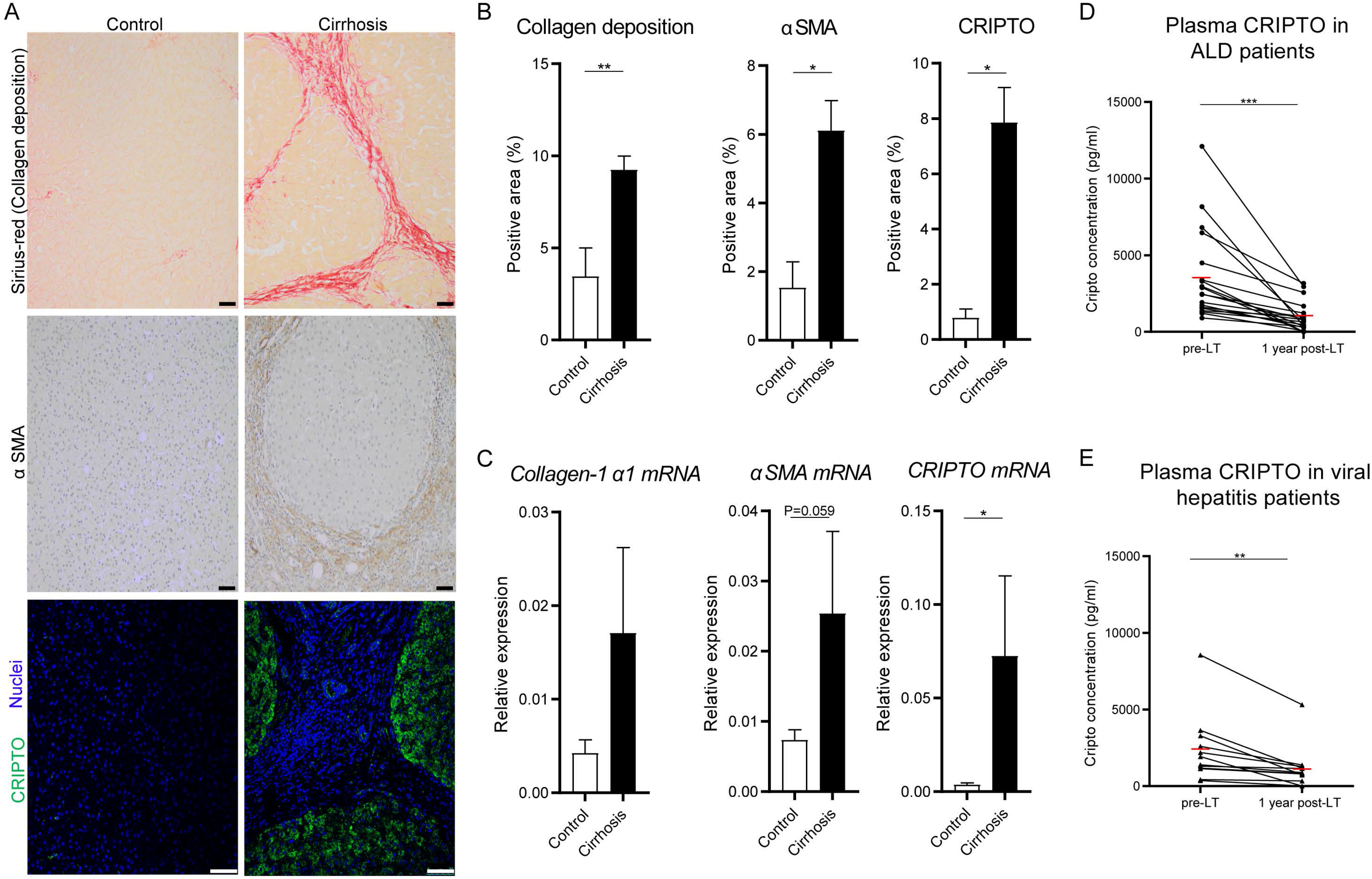
CRIPTO expression in patients with end-stage liver cirrhosis. Liver tissue samples of patients with ALD or viral induced liver cirrhosis (N=19) and controls (N=5) were randomly selected to evaluate CRIPTO expression. (A) Representative pictures of control and cirrhotic liver tissue stained for collagen deposition (Sirius-Red, scale bars 50um), αSMA (scale bars 50um) and CRIPTO (green), Nuclei stained with DAPI (Blue) scale bars 75um). (B) Quantification of Sirius-red, αSMA and CRIPTO staining (mean±SEM). (C) mRNA expression levels of *Collagen-1*α*1*, α*SMA* and *CRIPTO* normalized to β-actin (mean±SEM). *p≤0.05 **p≤0.01. (D-E) CRIPTO levels in plasma decrease after liver transplantation (LT) in different aetiological sub-cohorts. CRIPTO levels in pre- and post-LT paired plasma samples of patients suffering from (D) ALD (N=19) or (E) viral (N=12) induced liver cirrhosis. Mean group levels are indicated by a red line. **p≤0.01 ***p≤0.001.

**Figure 2.**
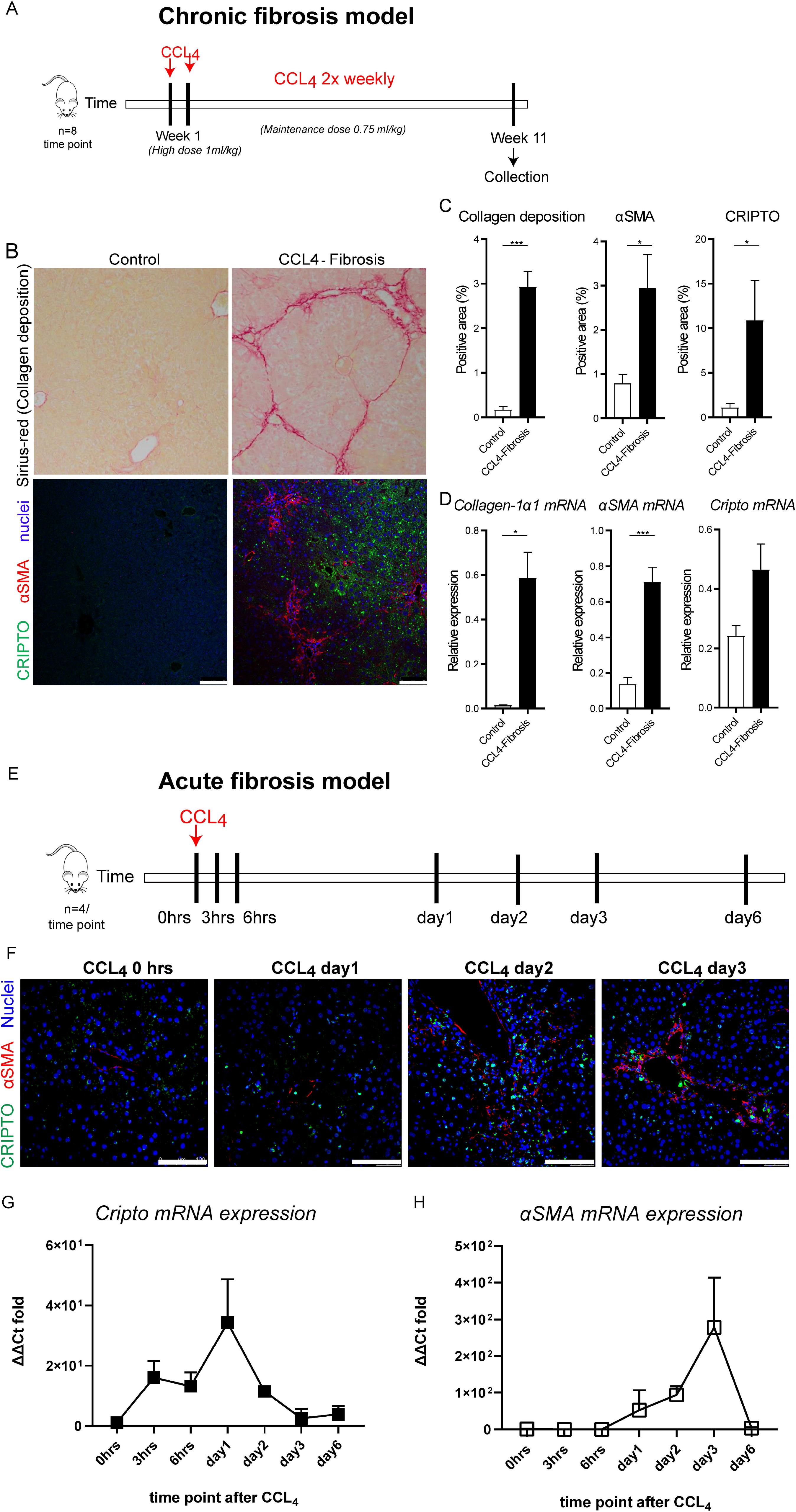
CRIPTO is upregulated in a CCL_4_ mouse model for chronic liver fibrosis and is reactivated in the early phase of acute CCL_4_ liver fibrosis. (A) Experimental setup. Mice received chronic administration (11 weeks) of CCL_4_ to induce liver fibrosis. (B) Representative pictures of healthy control and fibrogenic liver tissue stained for collagen deposition (Sirius-Red), and co-labeled for αSMA (red) and CRIPTO (green) (scale bars 75um). (C) Quantification of Sirius-Red, αSMA and CRIPTO staining (N=8 mice, mean±SEM). (D) mRNA expression levels of *collagen-1*α*1*, α*SMA* and *Cripto* normalized to β-actin (N=8 mice, mean±SEM). *p≤0.05 ***p≤0.001 (E) Experimental setup. Mice received one shot of CCL_4_ (acute liver fibrosis model, 1mg/kg) and liver tissues were collected at day 0; 0, 3, 6 hours, day 1, day 2, day 3 and day 6 (n=4 per time point). (F) Representative images of fibrotic liver tissues stained for αSMA (red) and CRIPTO (green) at day 0, day 1, day 2 and day 3. Nuclei are stained with TO-PRO-3. Scale bars 100um. (G) mRNA expression levels of *Cripto* and (H) α*SMA* normalized to *Gapdh* (Normalized ΔΔCt fold change to timepoint 0 hrs, mean±SD).

### RNA isolation, cDNA synthesis and quantitative Polymerase Chain Reaction (qPCR)

Mouse and human liver tissues were homogenised with UltraTurrax homogenizer (T25 basic, IKA) and TRIpure reagent (Roche). Subsequently, mRNA was isolated following TRIpure RNA isolation protocol. Promega standard protocol was used to synthesise cDNA from 1 μg RNA (Promega, Madison, Wisconsin, USA). Cripto, collagen-1α1 and αSMA expression in human and mouse samples were measured by qPCR analysis. qPCR reaction mixtures consisted of 5 μl iQ SYBR Green supermix reagent (Bio-Rad Laboratories, Berkeley, California, USA, 1708886), 1 nM primers and 4 μl cDNA. Results were normalised to β-actin for mouse and human samples. The primer sequences are indicated in **Supplementary table 2**.

### Cell culture and reporter assay

Human cell lines 293T and hepatic stellate cell lines LX-2 were maintained in Dulbecco`s Modified Eagle Media (DMEM) supplemented with 10% fetal calf serum and 1% penicillin/streptomycin. For assessing αSMA expression, 293T were serum starved overnight and stimulated with TGF-β3 5ng/ml, recombinant Cripto protein or transfected with Cripto expression plasmid (pcCripto) using Lipofectamine2000 (Life Technologies, Carlsbad, USA) according to the manufacturer`s instructions. For reporter assay, 293T cells were seeded at a density of 50.000 cells in 500ul medium in 24-well plates and transfected with 100ng αSMA reporter αSMA-luc plasmid, 10ng CAGGS-Renilla luciferase. The Firefly luciferase and Renilla luciferase levels in the lysates were measured using Dual Luciferase Assay (Promega, Madison, USA). For overexpression by adenoviral mediated delivery, 200 multiplicity of infection (MOI) of high titer virus was incubated with the media for 24 hours.

### Plasma Cripto measurements

Cripto levels in human plasma samples were measured using ELISA, performed according to manufacturer’s protocol (R&D systems, Minneapolis, Canada, DY145).

### Statistical Analysis

IBM SPSS statistics software (SPSS Inc. Chicago, IL USA, version 23) was used to perform Spearman tests for correlations. GraphPad Prism software (GraphPad Software, version 5.01, San Diego, CA) was used to perform Student’s t-test for the comparison between 2 groups. P-values lower than 0.05 were considered to be statistically significant. The data in the graphs are presented as means ± standard error of the mean (SEM).

## Results

### Cripto expression in patients with end-stage liver cirrhosis

The presence of liver fibrosis was evaluated by Sirius-red stained collagen-1α1 deposition and αSMA stained activated stellate cells. Liver tissue of patients with cirrhosis showed significantly higher Sirius-red and αSMA positive signal compared to control tissue, which confirmed the clinical diagnosis of cirrhosis (**Fig. 1A and B**). Cripto signal in these tissues was mainly observed in the hepatocytes and was clearly more present in the cirrhotic tissue as compared to control tissue, with 16 out 19 (84.2%) cirrhotic livers showing Cripto signal above the highest level in control livers (**Fig. 1A and B**). Furthermore, the results showed a positive correlation between the level of CRIPTO staining and the laboratory MELD scores of the patients (correlation coefficient: 0.577, P<0.003). No significant correlations between CRIPTO and Sirius-red or αSMA staining were observed (data not shown). QPCR analysis also showed elevated *collagen-1*α*1*, α*SMA* and *Cripto* mRNA expression levels in cirrhotic liver tissues compared to control liver tissues (**Fig. 1C**). Altogether, these results indicate that livers of patients with cirrhosis express higher levels of CRIPTO compared to control liver tissue and that the level of CRIPTO staining is correlated to the MELD score.

ELISAs were performed to study whether CRIPTO is reflected in blood of patients with liver cirrhosis. Cripto was detected in 31 out of 45 plasma samples of patients with end-stage cirrhosis (69%) and only in 2 out of the 16 controls (13%) (Chi-square 15.1; p<0.001). The mean Cripto level was significantly (p=0.03) higher in the end-stage liver cirrhosis group compared to that of the healthy controls (**Supplementary Table 1**). Circulating CRIPTO levels did not correlate with the MELD score (correlation coefficient: 0.151, p=0.310).

One year after LT, Cripto levels had significant decreased in all 31 patients as compared to their initial level before transplantation (**Supplementary Table 1, Fig. 1D and E**). A significant decrease in plasma CRIPTO concentration was also observed when the ALD and viral-induced cirrhosis cohorts were analysed separately (**Fig. 1D and E**). Altogether these data indicate that Cripto level in plasma decreases significantly once the cirrhotic liver has been replaced by a healthy donor liver. Nevertheless, the post-LT plasma Cripto levels still remained considerably higher than in the healthy controls, though not statistically significant at the Cripto level (p=0.6) but very clear at the positive frequency level (detectable Cripto level in patients 27/31 versus 2/16 for controls, Chi-square 24.9; p<0.0001).

### Cripto expression in patients with end-stage heart failure

The presence of cardiac fibrosis was evaluated by the expression of collagen I and aSMA-staining was used to visualize the MFBs. Clear collagen I –positive areas were found in myocardial samples of heart failure patients (**Supplementary Fig. 1**). Cripto expression, however, was hardly detected. Limited expression was observed in the interstitial cells, which did not express αSMA. Also areas with high density of MFBs did hardly show Cripto expression (not shown). These observations show that, in contrast to liver fibrosis, Cripto is only limitedly expressed in end-stage heart failure patients.

### Cripto expression in mouse models of liver fibrosis

To further study the potential role of Cripto in liver fibrosis we evaluated Cripto expression in a CCl_4_-induced mouse model for chronic liver fibrosis (11 week induction; **Fig. 2A**). Liver fibrosis was confirmed by Sirius-red and αSMA staining of the paraffin embedded liver tissue (**Fig. 2B**). Quantification of the Sirius-red and αSMA staining revealed higher content of collagen deposition and activated stellate cells in the livers of mice with fibrosis compared to healthy control animals (**Fig. 2C**). Similar to the observations from the human clinical data (**Fig. 1**), Cripto staining was pronounced in the liver tissues of mice with fibrosis and mainly observed in the hepatocytes (**Fig. 2B and C**). These findings were further supported by qPCR analysis, which also showed higher *collagen-1*α*1*, α*SMA* and *Cripto* RNA expression in the livers of mice with liver fibrosis compared to the healthy control livers (**Fig. 2D**).

To study the kinetics, dynamics and stimulus of the reactivated Cripto expression, we used an acute liver fibrogenesis model in which mice were subjected to a single shot of CCl_4_ and subsequently analysed liver tissues at various time points (**Fig. 2E**). Immunofluorescent co-staining of Cripto and αSMA indicated the presence of rare Cripto positive cells in the central vein area, as early as 24 hours post CCl_4_ shot (**Fig. 2F**; day1) followed by a short term expansion during day 2 when the MFBs (αSMA expressing cells) start to accumulate. αSMA protein expression levels reach there maximum at day 3 after CCl_4_ administration. Within these first 3 days only few Cripto-positive cells were observed (**Fig. 2F**; day 3). At the mRNA level, Cripto was upregulated at 3 hours after CCl_4_ exposure, reached highest expression on day 1 and rapidly reversed to low levels already on day 3 (**Fig. 2G**). Instead, αSMA mRNA levels increased later on (day 2) and reached maximum level on day3 (**Fig. 2H**). These results might suggest that Cripto expression is silenced in normal liver tissues, and is induced as an immediate response to cell injury and preceded activation of αSMA-positive MFBs. In this model, complete regeneration of the liver and resolution of fibrosis around the damaged portal triads, occured within seven days from the CCl_4_ administration **(Fig. 2G-H)**.

### Cripto expression in mouse models of cardiac fibrosis

To study the potential involvement of Cripto in fibrosis further, its expression was also determined in 3 different mouse models of cardiac fibrosis i.e. MI, pressure overload and an *ex vivo* fibrosis model for whole mouse hearts. Cripto was virtually absent in healthy non-operated (**Fig. 3A**) and in sham-operated mouse hearts (**Fig. 3L**). However, upon MI induction by permanent ligation of the left anterior descending artery, a progressive increase in the expression of Cripto was observed. At day 1 after MI, single Cripto-positive interstitial cells were observed (**Fig. 3B**). Day 3 after MI, Cripto became expressed at the infarction side of the border zone of the infarcted heart, opposite to the αSMA-expressing cells, which were present at the non-infarction side of the border zone (**Fig. 3C and D**). At day 7 after MI, Cripto and αSMA were expressed throughout the infarcted area (**Fig. 3E**). At day 14 after MI, the expression of both Cripto and αSMA had diminished near the border zone (**Fig. 3F**) and was only present in the center of the infarcted area (**Fig. 3F and G**). At 28 days after MI, both Cripto and αSMA had diminished in the infarcted heart (**Fig. 3H**). Pressure overload of the left ventricle was induced by transverse aortic constriction. After 2 weeks, upregulation of Cripto and αSMA expression was observed surrounding the coronary arteries (**Fig. 3I and K**), which had decreased 6 weeks after pressure overload, whereas increased collagen I expression remained (**Fig. 3J and N**), indicating perivascular fibrosis. At 8 weeks after pressure overload, upregulation of Cripto, αSMA and collagen I was observed in between the cardiomyocytes of the ventricular wall (**Fig. 3Q-S**), indicating interstitial fibrosis. Culturing of whole mouse hearts in the miniature tissue culture system (MTCS) under specific conditions resulted in myocardial fibrosis. Staining for Cripto indicated that the formation of αSMA-positive cells coincides with increased Cripto staining (**Fig. 3T-V**). Overall these observations show a clear association of cardiac fibrosis and temporary Cripto expression.

**Figure 3.**
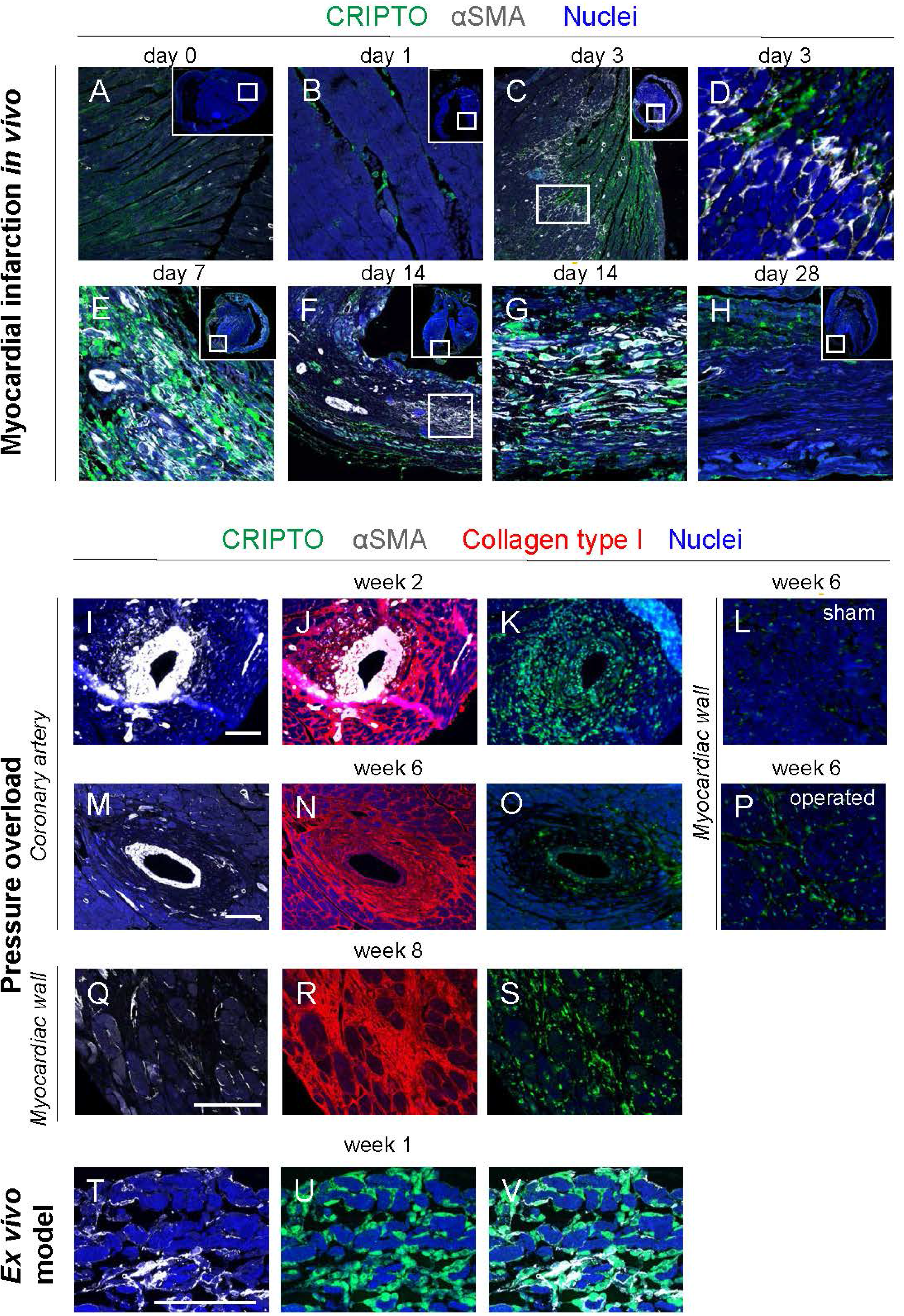
CRIPTO reactivation during wound healing response to cardiac tissue injury. CRIPTO (green) expression during myocardial infarction associated with αSMA expression (white) and/or collagen type I (red) expression during myocardial infarction (A-H; A: day 0, B: day 1, C-D: day 3, E: day 7, F-G: day 14, H: day 28), pressure overload (I-K: 2 weeks, L-P: 6 weeks, L=sham, Q-S: 8 weeks); around coronary artery (I-O) and myocardiac wall (L, P-S), and during culture in the *ex vivo* cardiac fibrosis model (T-V: 1 week).

### Adenoviral-mediated *in vivo* overexpression of Cripto leads to increased fibrogenesis

To identify whether the observed high Cripto levels play a causal role in the induction of wound healing, we combined the acute CCl_4_ model with *in vivo* overexpression of Cripto. Adenovirus carrying Cripto-coding sequence (AdCripto) or control adenoviral particles (AdlacZ) were injected intravenously for optimum uptake by the liver, 24 hours prior to the administration of CCl_4_. Liver tissues were analysed at day 1, 3 and 6 after CCl_4_ (**Fig. 4A**). Expression of αSMA and collagen type I deposition was highest at day 3 (**Fig. 4B and C**) after CCl_4_, and was higher in the AdCripto group compared to control groups. Cripto expressing cells were consistently observed after fibrogenic induction by CCl_4_ in both groups (AdlacZ+CCl_4_ and AdCripto+CCl_4_) (**Fig. 4C**). Assessment of the mRNA levels of αSMA and Collagen type I chain (Col1A1), showed no upregulation after exogenous delivery of Cripto-coding sequence (AdCripto group), but only increased following CCl_4_ administration (**Fig. 4D and E**). Interestingly, Col1A1 mRNA levels were significantly increased in the combined AdCripto+CCL4 group on day 3, compared to all other treatment groups (**Fig. 4E**,* p◻0.05). Quantification of αSMA staining showed higher levels in the AdCripto+CCl_4_ (**Fig. 4F**) (**** p◻0.0001 vs AdlacZ, **** p◻0.0001 vs AdCripto, p=0.08 vs AdlacZ+CCl_4_). Quantification of Collagen staining showed similar trend of highest levels in the AdCripto+CCl_4_ group (**Fig. 4G**) (**** p◻0.0001 vs AdlacZ, **** p◻0.0001 vs AdCripto, p=0.18 vs AdlacZ+CCL4). Using *in vitro* assays we have identified Cripto as an upstream direct regulator of αSMA, marker of transdifferentated MFBs, the main cell type responsible for fibrogenesis (**Supplementary Fig. 2**). Transient overexpression of Cripto or treatment of 293T cells with recombinant Cripto protein upregulated αSMA protein expression (**Supplementary Fig. 2A**) and activated αSMA gene reporter to an equivalent manner as treatment with profibrotic TGFβ cytokine (**Supplementary Fig. 2B**). Similarly, stable overexpression of Cripto in LX2 human liver fibroblasts led to α*SMA* gene expression levels (**Supplementary Fig. 2C**).

**Figure 4.**
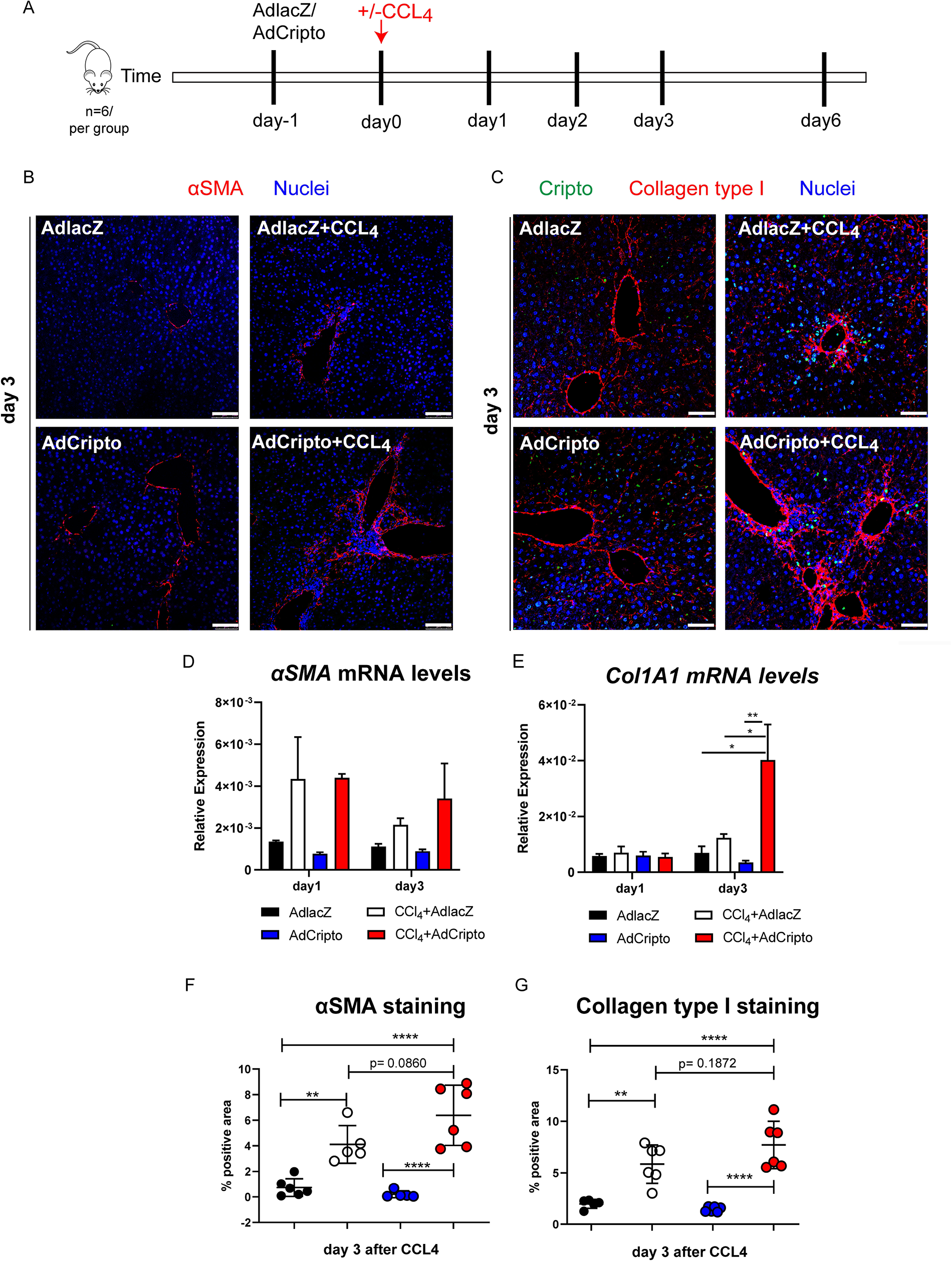
In vivo overexpression of CRIPTO in combination with CCL_4_ exacerbates fibrosis. (A) Experimental setup. Mice received Adenovirus expressing Cripto or lacZ (AdlacZ, AdCripto 10E+9 particles intravenously) 24 hours prior to one shot of CCL_4_ (acute liver fibrosis model, 1mg/kg) and liver tissues were collected at day 1, day 2, day 3 and day 6 (n=6 per treatment group; AdCripto+/− CCL_4_, AdlacZ +/− CCL_4_). (B) Representative images of fibrotic liver tissues stained for αSMA (red) on day 3 after CCL_4_. Nuclei are stained with DAPI. Scale bars 100um (C). Representative images of fibrotic liver tissues stained for Cripto (green), Collagen type I (red) on day 3 after CCL_4_. Nuclei are stained with DAPI. Scale bars 50um. (D-E) mRNA expression levels of α*SMA* and *Col1a1* normalized to *actin*, n=2 per time point and per group. Ordinary two-way ANOVA, *p≤0.05, **p≤0.01. (F-G) Quantification of αSMA and CRIPTO staining (average n=3 fields of view per tissue; n=2 mice, mean±SD). Ordinary two-way ANOVA, **p≤0.01, ****p≤0.0001.

Overall, Cripto upregulation increased the relative level of fibrosis as showed by more accumulating MFBs and higher expression of fibrotic markers *in vivo* and *in vitro*.

### Functional inhibition of Cripto by ALK4Fc ligand trap reduces collagen deposition and stimulates hepatocyte proliferation

To further understand the dynamics and functional role of Cripto in the context of fibrogenesis, we inhibited Cripto-secreted protein form to sequester the Cripto-expressing cells during fibrotic induction in the acute liver CCl_4_ fibrosis model. The ligand trap for the extracellular binding domain of ALK_4_ receptor, binding partner of secreted Cripto (ALK4Fc) [44, 45] was administered i.p. *in vivo*, 2_4_ hours prior to and at day 1 and 2 after CCl_4_ exposure (**Fig. 5A**). Liver tissues were analysed at day 1 and day 3 after CCl_4_ administration, for Cripto, proliferation (Ki67) and fibrotic (SMA, Collagen type-I, −III) markers. Cripto-positive cells appeared around the damaged area (indicated by Collagen type-I, **Fig. 5B** in red) in both conditions (CCl_4_+IgG, CCl_4_+ALK4Fc) at day 1 and decreased at day 3 (**Fig. 5B**). Proliferating Ki67-positive hepatocytes were apparent outside the fibrotic areas on day 3 (**Fig. 5C**). Furthermore, significantly higher levels of Ki67-positive hepatocytes were observed in the CCl_4_+ALK4Fc liver tissues, compared to the CCl_4_+IgC tissues (**Fig. 5E**, * p◻0.05). Collagen type I staining was significantly lower in CCl_4_+ALK4Fc liver tissues (**Fig. 5C**) compared to CCl_4_+IgC (**Fig. 5F**, day3, *** p◻0.001). αSMA expression indicated smaller fibrotic areas around the central veins of the ALK4FC group, day 3 (**Fig. 5D and G**, ns, p=0.09). Similarly, collagen type III deposition was decreased in the CCl_4_+ALK4Fc group (**Fig. 5D and H**, * p◻0.05). Overall, the data showed decreased levels of all fibrotic markers and increased levels of regenerating hepatocytes in livers exposed to a combination of CCl_4_ and Cripto antagonist (ALK4FC).

**Figure 5.**
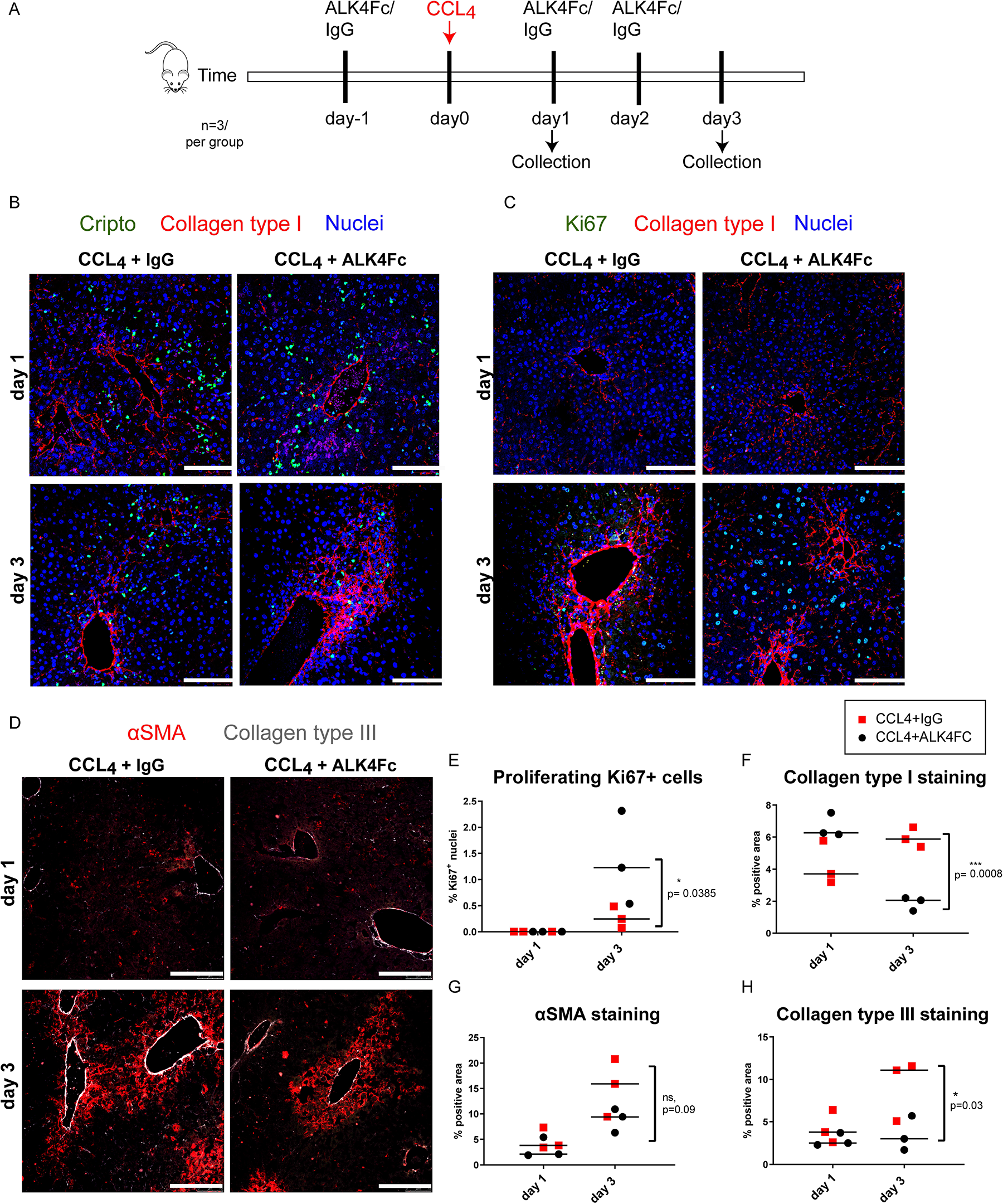
Blocking CRIPTO function by ALK4Fc leads to reduced collagen deposition in the acute CCL_4_ fibrosis model. (A) Experimental setup. Mice received ALK4Fc or IgG vehicle (5mg/kg) 24 hours prior to one shot of CCL_4_ (acute liver fibrosis model, 1mg/kg), and once daily for 3 days following CCL_4_. Liver tissues were collected at day 1 and day 3 (n=3 per treatment group; ALK4Fc+ CCL_4_, IgG+ CCL_4_). (B) Representative images of fibrotic liver tissues stained for Cripto (green) and Collagen type I (red) on day 1 and 3 after CCL_4_. Nuclei stained by DAPI. Scale bars 100um. (C) Representative images of fibrotic liver tissues stained for Ki67 (green) and Collagen type I (red) on day 1 and 3 after CCL_4_. Nuclei stained by DAPI. Scale bars 100um. (D) Representative images of fibrotic liver tissues stained for αSMA (red) and Collagen type III (gray) on day 1 and 3 after CCL_4_. Scale bars 100um. (E) Quantification of proliferating Ki67+ cells (percentage over the DAPI stained nuclei) (p=0.0385 *, CCL_4_+ALK4Fc vs CCL_4_+IgG day 3, two-way ANOVA, n=3). (F) Quantification of Collagen type I staining (p=0.0008 ***, CCL_4_+ALK4Fc vs CCL_4_+IgG day 3, two-way ANOVA, n=3). (G) Quantification of αSMA staining (n=3, mean) Ordinary two-way ANOVA, p=0.09 CCL_4_+ALK4Fc vs CCL_4_+IgG day 3. (H) Quantification of Collagen type III staining (n=3, mean). Ordinary two-way ANOVA, p=0.03 *, CCL_4_+ALK4Fc vs CCL_4_+IgG day 3; *p≤0.05 ***p≤0.001.

## Discussion

Fibrosis may manifest in multiple tissues such as skin, lung, heart, kidney and liver [46], while antifibrotic treatments directly targeting the process of fibrogenesis are currently not available and activating mechanisms are not fully elucidated [47]. Excessive fibrosis develops on the basis of chronic inflammation and epithelial cell injury due to infections, environmental hazards or acute tissue damage. Fibrogenesis is a necessary part of the would healing response, while fibrosis is aberrant fibrogenesis (extracellular matrix deposition) which does not resolve or facilitate tissue regeneration. Some tissues, such as liver with high regenerative capacity (cell replenishment of the injured cell type), may reverse fibrostic processes if the tissue damaging stimulus is removed early in the disease development, Other organs such as the heart have a low regenerative capacity resulting in the permanent presence of fibrosis.

In the present study we assessed whether Cripto is expressed during fibrosis in multiple organs in humans and murine models. We observed Cripto expression in hepatocytes of human cirrhotic liver tissue, which is in line with the findings in a recent study where we showed that Cripto was highly expressed in HCC, and was associated with Sorafenib resistance [36]. The levels of Cripto expression in the hepatocytes of HCC patients positively correlate with the MELD scoring system for end-stage liver diseases [48, 49], illustrating that Cripto expression is related to the severity of the disease. In addition, we observed elevated Cripto levels in most (69%) of the measured plasma samples of patients with end-stage liver disease, which decreased after the patients underwent LT. This observation is in line with recent findings of Zhang et al. whom also observed enhanced Cripto levels in serum of patients with HCV- and HBV-induced cirrhosis [50]. In acute and chronic mouse models of liver fibrosis we also observed upregulation of Cripto indicative for a general and well preserved role of Cripto during fibrogenesis.

Whereas in the MI, pressure overload and ex vivo mouse models of cardiac fibrosis Cripto was upregulated, only a very limited amount of Cripto was observed in the human samples of cardiac fibrosis. The difference can likely be explained by the dynamic expression of Cripto during the different stages of fibrosis. The samples were derived from patients that suffered from end-stage heart failure suggesting the long existence of fibrosis in the heart. In a recent study Cripto was detected in blood samples of patients with planned MI within 1 hour [51], suggestion that Cripto is expressed shortly after cardiac injury. Together with our observation in the MI model, these observations suggest that Cripto is mostly present in the early phases of fibrosis.

Although in most plasma samples of patients with end-stage liver cirrhosis elevated Cripto levels were observed, in about 30% Cripto was undetectable, which is in concordance with a previous study [50]. This might indicate that also in chronic liver fibrosis Cripto expression is dynamic.

The Cripto-positive cells in the acute CCl_4_ liver model accumulated in the damaged areas and only partially coinciding with MFBs, indicating that the source of Cripto could be a distinct cell type, possibly bone marrow fibrocyte, resident progenitors or hepatic stellate cells prior to their transdifferentiation to MFBs or recruited inflammatory cells [52]. In contrast, in advanced stages (human cirrhosis and chronic mouse liver fibrosis model), Cripto is predominantly expressed by epithelial cells (hepatocytes) surrounding but not within the fibrotic areas indicating that the cellular source of Cripto is different in advanced progression, and derives from intrinsic cell alterations in damaged hepatocytes. In the injured heart, the expression of Cripto is initially, similar to liver, present in interstitial non-MFBs. Shortly thereafter, Cripto expression mostly coincides with the expanding population of MFBs until fibrotic area has been established after which the Cripto expression ceases. Expression of Cripto in the cardiomyocytes however is virtually absent. Ex vivo induced cardiac fibrosis similarly led to the appearance of Cripto-positive cells, which are therefore tissue resident and not derived from a systemic source.

The difference in cell type in which Cripto is expressed between organs is intriguing. Whereas in regenerating tissues like the liver and skeletal muscle, Cripto is expressed in the progenitor cells of the newly formed parenchymal cells, in the non-regenerating heart Cripto remains expressed in the stroma cells. During cardiac development, Cripto is required for cardiomyogenesis [53–56]. The lack of reactivation of Cripto in the cardiomyocytes could be hypothesized to preclude myocardial regeneration.

Adenoviral Cripto overexpression in the acute CCl_4_ liver mode resulted in higher levels of fibrotic markers, whereas inhibition of Cripto by the ALK4Fc ligand trap [45], led to improved hepatocyte proliferation and significantly reduced fibrosis. These observations indicate that Cripto is not simply a mere consequence of the tissue damage or fibrosis, but it directly regulates the cascade of fibrosis by inducing myofibroblast formation and collagen production. The expression of Cripto in the heart shortly after MI concomitant with, and even preceding, the appearance of MFBs, suggests a similar role for Cripto in cardiac fibrosis.

From a molecular point of view, the mechanism causing the upregulation of Cripto and the enhanced fibrotic response remains to be further addressed. NANOG, a regulator of Cripto expression, is expressed in hepatocytes during fibrogenesis [36, 57], and expressed in cardiac interstitial cells [58], which could thus contribute to the Cripto expression during fibrogenesis [59]. Furthermore, Cripto expression has been shown to be involved in the activation of the well-known SMAD pathways, which could promote fibrogenesis and eventually HCC formation [33, 57, 60] and regulate the cardiac fibrotic response [61]. It has been shown that inducing cell damage to HepG2 cells leads to the upregulation of Cripto which initiates apoptotic resistance and increased proliferation via NF-κB/Survivin pathways [62]. A similar mode of Cripto reactivation may occur in vivo, in which Cripto may become re-expressed as a response to cellular injury in order to promote tissue regeneration by orchestrating the reactivation of both fibrogenic cells and of quiescent hepatocytes.

Cripto has a profibrotic role in both heart and liver fibrosis and we show for the first time that Cripto reactivation is linked to tissue homeostasis, wound healing response and fibrosis, and is conserved in different tissues with low and high regenerative capacity.

Altogether the observations from this study warrant further research to disentangle whether Cripto has a functionally relevant role in fibrotic diseases, with biomarker and therapeutic value while targeting its activation can alleviate the extent and progression of fibrosis.

## Acknowledgements

We thank Peter C.Gray for providing materials and revising the manuscript, Midory Thorikay for the adenoviral constructs, Kirsten Lodder and Tessa van Herwaarden for their expert technical assistance.

## Author Contributions

Conceptualization, Boudewijn Kruithof and Marianna Kruithof de Julio; Formal analysis, Sofia Karkampouna and Danny van der Helm; Investigation, Sofia Karkampouna, Danny van der Helm, Boudewijn Kruithof and Marianna Kruithof de Julio; Methodology, Boudewijn Kruithof and Marianna Kruithof de Julio; Resources, Bart Van Hoek and Marie José Goumans; Visualization, Hein Verspaget and M.J. Coenraad; Writing – original draft, Sofia Karkampouna, Boudewijn Kruithof and Marianna Kruithof de Julio; Writing – review & editing, Danny van der Helm, Bart Van Hoek, Hein Verspaget, Marie José Goumans and M.J. Coenraad.

---

**Supplementary Figure 1.**
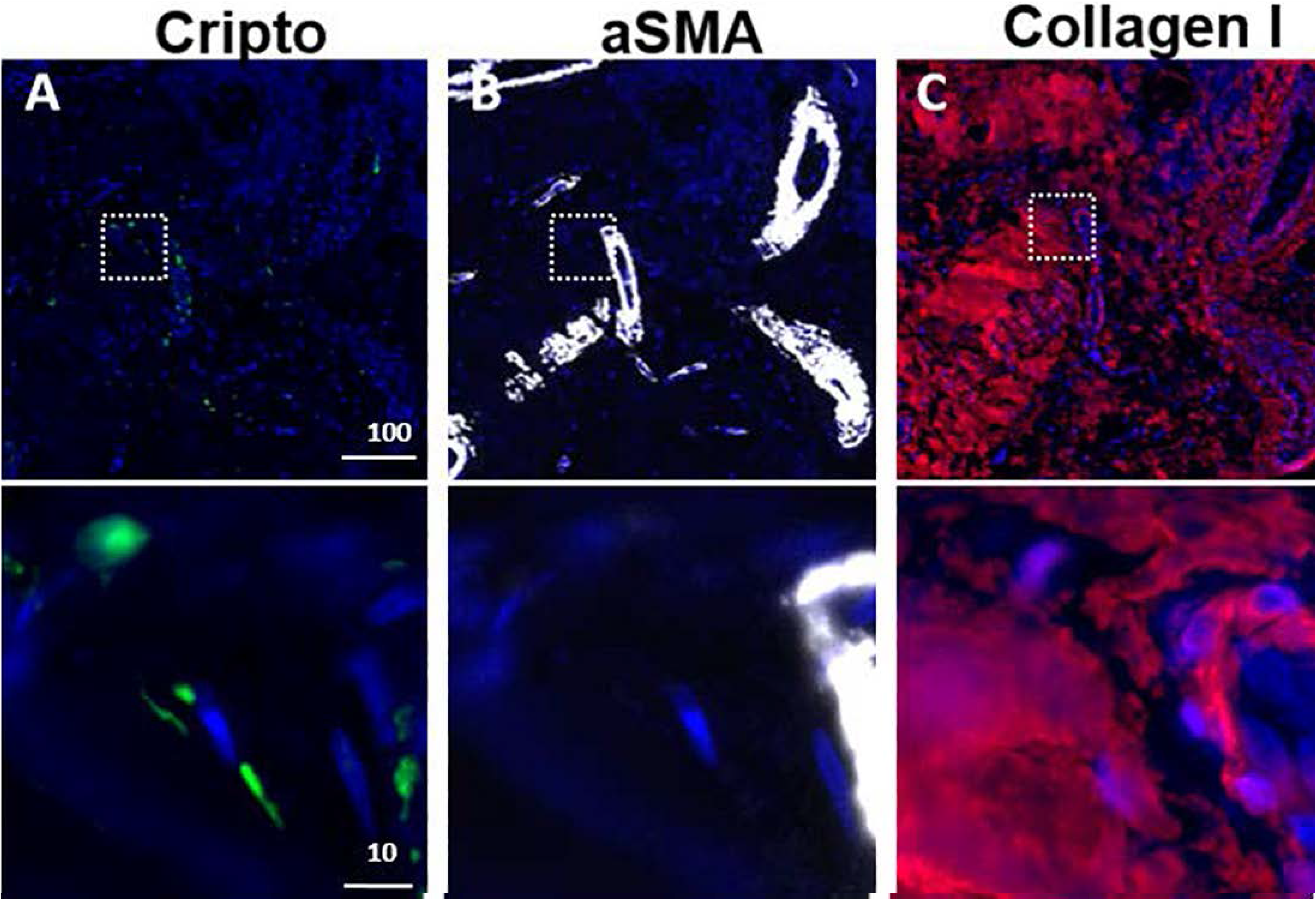
Cripto expression in patients with end-stage heart failure. Cardiac tissue samples of patients with end-stage heart failure (n=5) were stained for Cripto (A) aSMA (B) and collagen I (C). Scale bar 100 um.

**Supplementary Figure 2.**
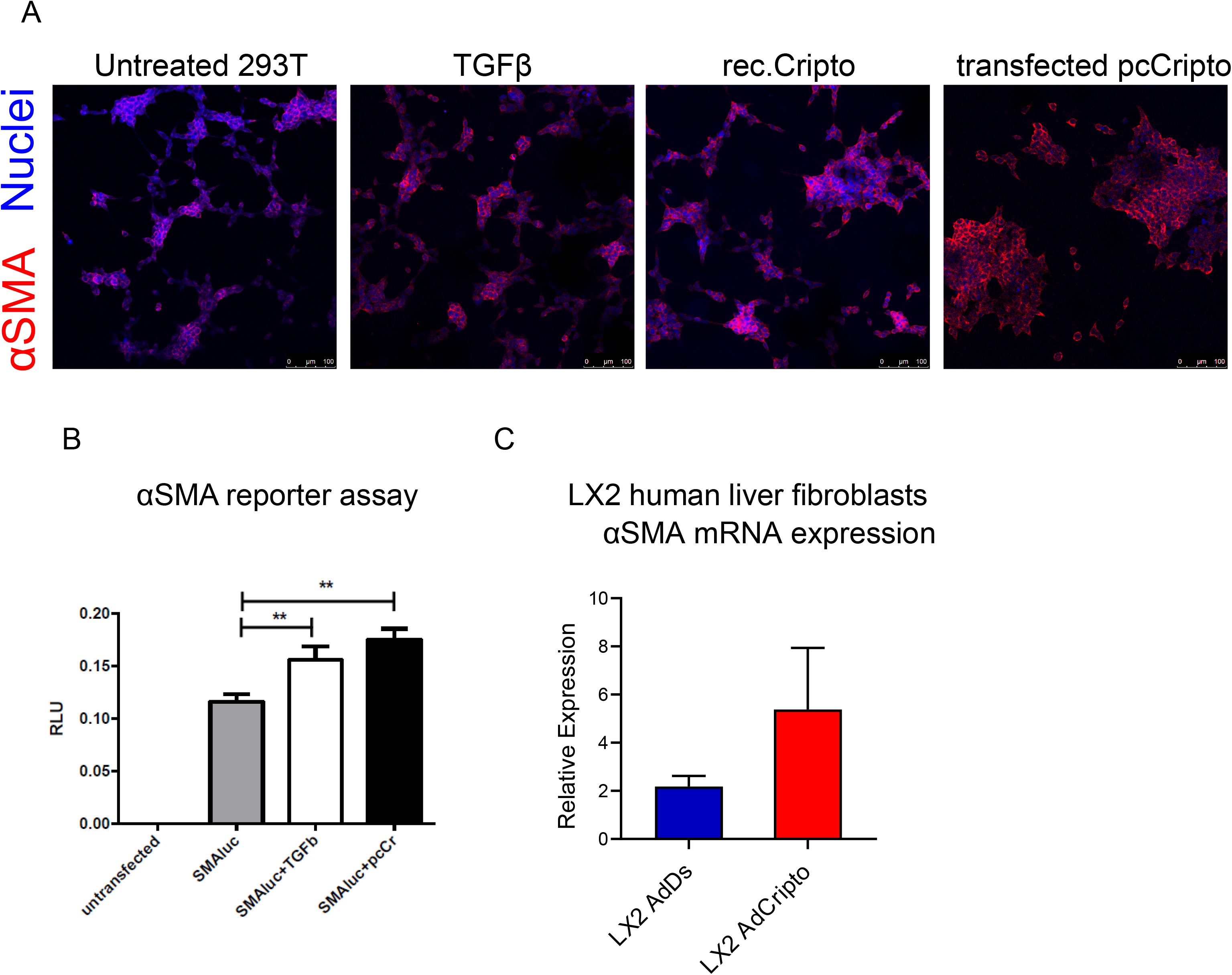
CRIPTO is an upstream regulator of αSMA expression. (A) *In vitro* effects of CRIPTO in 293 T cells either by transfection of full length CRIPTO sequence (pcCRIPTO) or by stimulation with recombinant CRIPTO protein. Cells were stimulated with TGF-β as positive control for αSMA induction. Cell staining with αSMA (red) and DAPI (nuclei). Scale bars 100um. (B) αSMAluc reporter assay on 293T cells, stimulated with TGF-β or transfected with CRIPTO-expressing plasmid (pcCRIPTO). (C) α*SMA* mRNA expression in LX2 human liver fibroblast cell line after adenoviral-mediated CRIPTO (AdCRIPTO) overexpression in vitro. AdDs; control adenovirus containing dsRED fluorescent protein.

**Supplementary table 1:** Patient characteristics. Data presented as median (range) for continuous variables and percentage (number) for categorized variables.

| Variable | Healthy controls<br>(N=16) | Pre-LT<br>(N=45) | Post-LT<br>(N=45) |
| --- | --- | --- | --- |
| Gender (male), % (n) | 50% (8) | 78% (35) |  |
| Age (median, range) | 29 (23-65) | 5<sub>4</sub> (42-69) |  |
| Aetiology |  |  |  |
| - Alcoholic liver disease |  | 25 |  |
| - Viral Hepatitis |  | 20 |  |
| Blood (median, range) |  |  |  |
| - AST (U/L) |  | 72 (2<sub>4</sub>-517) | 27 (11-240) |
| - ALT (U/L) |  | 37 (15-360) | 25 (8-401) |
| - INR |  | 1.2 (1-2.4) | 1.0 (0.9-2.4) |
| - ALP (U/L) |  | 130 (50-555) | 88 (47-487) |
| - Creatinin (μmol/L) |  | 92 (3<sub>4</sub>-171) | 111 (68-20<sub>4</sub>) |
| - γGT (U/L) |  | 43 (7-37<sub>4</sub>) | 39 (9-1395) |
| - Sodium (mmol/L) |  | 138 (12<sub>4</sub>-156) | 142 (13<sub>4</sub>-148) |
| - Bilirubin (μmol/L) |  | 46 (5-593) | 12 (5-29) |
| - Platelet count (10<sup>9</sup>/L) |  | 72 (30-142) | 14<sub>4</sub> (93-243) |
| Cripto plasma (pg/ml) | 0 (0-818) | 1381 (0-12108) | 357 (0-531<sub>4</sub>) |
| Clinical scores |  |  |  |
| - MELD |  | 15 (8-33) | 10 (6-18) |

LT = Liver Transplantation, ALD = Alcoholic Liver Disease, AST = Aspartate aminotransferase, ALT = Alanine aminotransferase, INR = International Normalized Ratio, ALP = Alkaline phosphatase, γGT: gamma-glutamyltransferase, MELD = Model for End-Stage Liver Disease

**Supplementary table 2:**
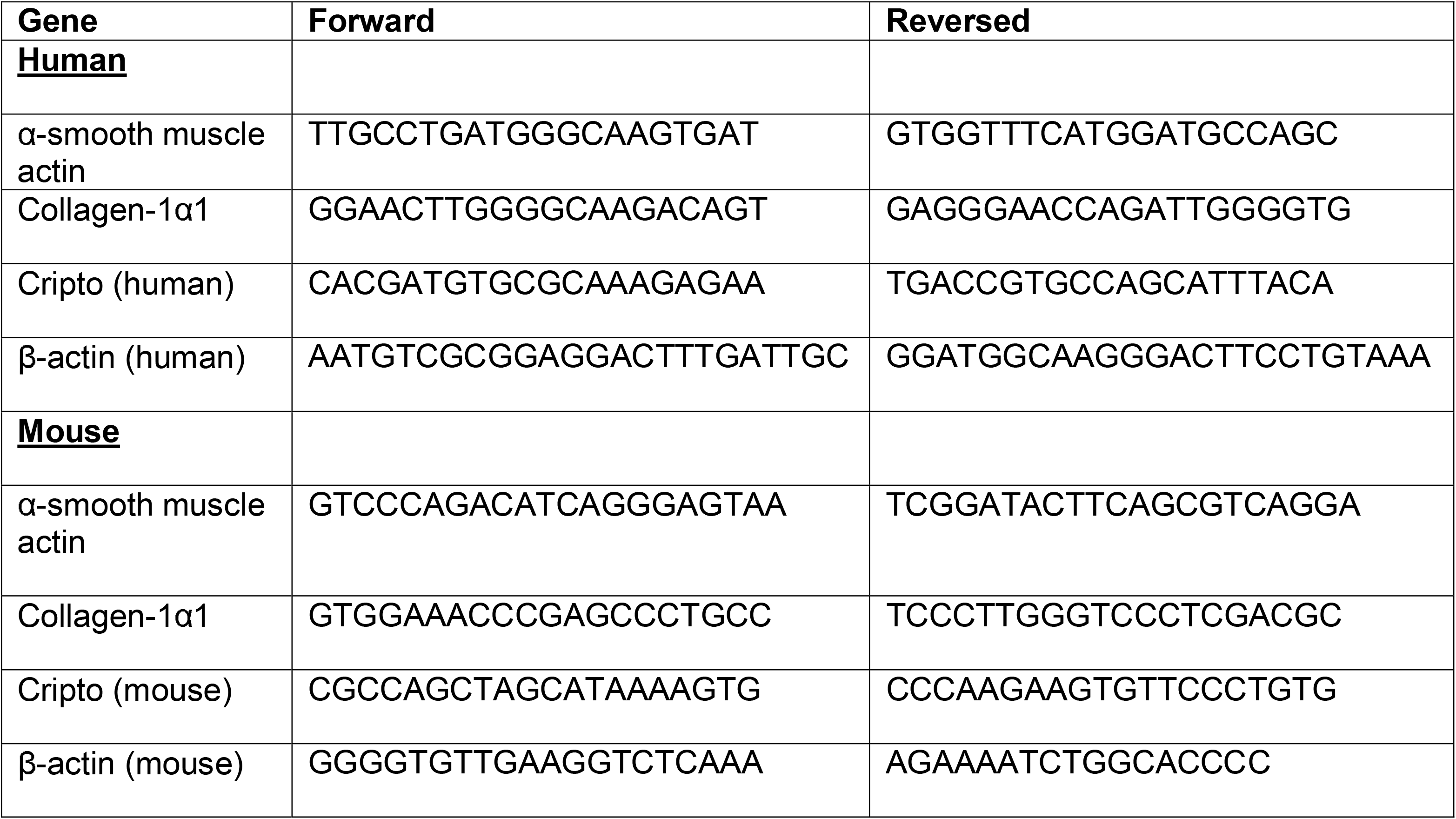
Primer sequences.

